# Tetrazoles as PPARγ ligands: A Structural and Computational Investigation

**DOI:** 10.1101/2021.02.17.431624

**Authors:** Karina de Paula, Jademilson C. Santos, Ana Carolina Mafud, Alessandro S. Nascimento

## Abstract

Diabetes is an important chronic disease affecting about 10% of the adult population in the US and over 420 million people worldwide, resulting in 1.6 million deaths every year, according to the World Health Organization. The most common type of the disease, type 2 diabetes, can be pharmacologically managed using oral hypoglycemic agents or thiazolidinediones (TZDs), such as pioglitazone, which act by activating the Peroxisome Proliferated-Activated Receptor γ. Despite their beneficial effects in diabetes treatment, TZDs like rosiglitazone and troglitazone were withdrawn due to safety reasons, creating a void in the pharmacological options for the treatment of this important disease. Here, we explored a structure-based approach in the screening for new chemical probes for a deeper investigation of the effects of PPARγ activation. A class of tetrazole compounds was identified and the compounds named T1, T2 and T3 were purchased and evaluated for their ability to interact with the PPARγ ligand binding domain (LBD). The compounds were binders with micromolar range affinity, as determined by their IC_50_ values. A Monte Carlo simulation of the compound T2 revealed that the tetrazole ring makes favorable interaction with the polar arm of the receptor binding pocket. Finally, the crystal structure of the PPARγ-LBD-T2 complex was solved at 2.3 Å, confirming the binding mode for this compound. The structure also revealed that, when the helix H12 is mispositioned, an alternative binding conformation is observed for the ligand suggesting an H12-dependent binding conformation for the tetrazole compound.

## 1. Introduction

The increasing prevalence of diabetes mellitus around the world raises important issues about the pharmacological treatment of this pathology. The Center for Disease Control and Prevention in the United States reported the prevalence of diabetes among the American population as about 10%, resulting in a burden of around US$ 327 billion (www.cdc.gov/diabetes). The management of the disease usually involves (*i*) the change in habits (nutritional restrictions and physical activity); (*ii*) the use of oral hypoglycemic agents (biguanides, sulfonylureas, or α-glucosidase inhibitors) and (*iii*) the use of insulin sensitizers, such as the thiazolidinedione (TZD) pioglitazone [1].

The TZDs act by activating the nuclear receptor Peroxisome Proliferator-Activated Receptor γ (PPARγ). This receptor is probably the most functionally diverse among the three PPARs, modulating a large repertoire of genes involved in glucose metabolism, lipid metabolism and energy dissipation, inflammation, among other functions [2]. Structurally, the PPARs are folded in the three domains typical for the nuclear receptor superfamily [3], with a short N-terminal domain, probably disordered; a central DNA binding domain folded as a zinc finger and a C-terminal ligand binding domain, folded as an alpha helix sandwich and that is responsible for ligand recognition, receptor dimerization and co-activators/co-repressors recruitment [4].

TZDs fully activate PPARγ leading to insulin sensitization and lowering the glucose levels [5]. Rosiglitazone and pioglitazone became important medicines in the treatment of type 2 diabetes mellitus, resulting in significant improvement in chronic hyperglycemia. However, posterior clinical findings showed that the use of rosiglitazone was associated with an increased risk of cardiovascular events [6,7], leading regulatory agencies around the world to restrict [8] or to suspend the use of rosiglitazone [9]. Curiously, pioglitazone was not shown to be associated with increased cardiovascular events to the same extent as rosiglitazone [10,11], suggesting that small changes in the stabilization of the ligand binding domain may reflect important differences in the pharmacological profile, as previously observed [12]. Bruning and coworkers showed that a range of binding modes is possible for PPARγ agonists, leading to different degrees of receptor activation and, possibly leading to a different spectrum of harmful effects [12,13].

PPARγ activation by TZDs is also known to promote adipocyte differentiation, leading to an increase in the number of small adipocytes in the tissue [14]. PPARγ is largely expressed in the adipose tissues and regulates the expression of several adipose target genes, such as lipoprotein lipase, fatty acid transporter protein, adipocyte fatty acid-binding protein aP2, acyl-CoA synthetase (ACS), phosphoenolpyruvate carboxy kinase (PEPCK), and malic enzyme, as reviewed elsewhere [15]. The *in vivo* effects of PPARγ activation on adipogenesis become clear in the studies of the receptor antagonists as the molecule SR-202, which inhibits adipose tissue accumulation and reduces body weight in wild type mice fed with a high-fat diet [16].

The experimental observation that PPARγ activation leads to a decrease in TNF-α levels raised the putative role of TZDs or synthetic ligands in the treatment of inflammation [5]. Several lines of evidence suggest that the anti-inflammatory effects of PPARγ activation may be useful for neuroprotection [17,18], management of endothelial function [19,20] and colitis [21]. Besides its action on TNF-α levels, the carrageenan mouse model suggests that PPARγ activation also regulates the level of myeloperoxidase, adenosine deaminase, IL-1, IL-17, and VEGF-α [22].

Although the beneficial effects of PPARγ activation can be clearly defined from its known molecular mechanism of action, safety issues related to the pharmacological therapy were observed with the TZDs. It was observed that full PPARγ agonists, such as the TZDs rosiglitazone and pioglitazone, cause more important cardiovascular effects than partial agonists or PPAR modulators, and the development of partial agonists became an ultimate goal in insulin sensitization mediated by PPARγ [23]. Larsen and coworkers showed that equi-efficacious doses of rosiglitazone and balaglitazone, a PPARγ partial agonist, resulted in the same hypoglycemic effect but with minor fluid retention and cardiac enlargement observed for the partial agonist [23,24]. The same effect has been confirmed for other PPARγ partial agonists [25–27].

Given the health and economic impact of diabetes mellitus, it is still important to develop new chemical entities (NCE) that could serve as insulin sensitizers acting through PPARγ. Those NCEs serve as chemical probes to show the impact of partial receptor agonism and its desired and side effects. Furthermore, the large number of crystal structures available nowadays for this receptor makes the structural-based methods for ligand development an attractive strategy for NCEs finding [28]. Here, we used a structure-based strategy to find a new scaffold of PPARγ partial agonists. A class of tetrazole compounds was identified and found to be PPARγ binders with an IC_50_ in the range of 5-20 μM. One compound of this class, T2, was co-crystallized with PPARγ, confirming its binding conformation.

## 2. Methods

### 2.1 Molecular docking and simulations

Crystal structures of PPARγ bound to rosiglitazone, PDB IDs 1FM6 [29] and 7AWC, were used for docking calculations. Polar hydrogens were added with UCSF Chimera [30] and charges were attributed to protein atoms according to AMBER FF14SB force field. The drugs-now subset of ZINC was used for molecular docking in UCSF DOCK 3.5 program [31].

After the first screening of docking, a subset of tetrazole compounds identified in the screen was used for further calculations using LiBELA [32]. The compounds were redocked into PPARγ-LBD crystal structure using an all-atom AMBER force field and obtained docked scores and docking poses were used in structural analysis. In this step, a modified version of the Energy Score previously described was used. Here, a softcore potential [32] was used together with a continuum electrostatic model [33], a 12-10 hydrogen bond potential, and a desolvation penalty [34]. The interaction energy (E_interact_) is thus described as shown in equation 1 below:

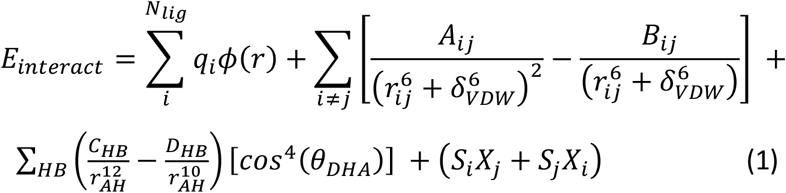

In equation (1), the interaction potential energy is described: the first term accounts for the polar interactions, and ϕ(r) is the electrostatic potential computed using a continuum electrostatic method with the program DelPhi [35–37]. The second term is a smoothed Lennard-Jones potential that accounts for the van der Waals interactions, with the smoothing constant δ_VDW_ set to 1.75 Å. *A*_*ij*_, *B*_*ij*_ and *r*_*ij*_ are the Lennard-Jones repulsive term, attractive term, and interatomic distance, respectively, taken from AMBER FF14SB force field. The third term is a 12-10 potential that accounts for the hydrogen bonds with a directionality term. Here r_AH_ is the distance between the hydrogen and the acceptor atom, C_HB_ and D_HB_ are constants derived to score an optimal HB at −5 kcal/mol and θ_DHA_ is the angle between donor, hydrogen and acceptor atom. The final term is the desolvation term, as previously described [34], with the constants α, β and σ set to 0.1 kcal/(mol·*e*^2^), −0.005 kcal/mol and 3.5 Å, respectively. The computer code implemented in LIBELA is open and available in https://github.com/alessandronascimento/LiBELa.

The docked poses were afterward used for Monte Carlo (MC) simulations using a modified version of LiBELa, which we named MCLiBELa. For these simulations, the receptor is kept rigid and is represented by pre-computed interaction potential grids. For each MC step, the ligand is randomly rotated and translated, and each rotatable bond is also randomly shifted. A *move* is then defined as a set of the variables Δx, Δy and Δz, defining a random shift in the ligand applied to its center of mass, and Δα, Δβ and Δγ, that define a random rotation in Euler angles around the ligand center of mass. Additionally, ligands with rotatable bonds will have n_rot_ Δθ variables, where n_rot_ is the number of rotatable bonds and Δθ denotes a random shift in each rotatable bond. The maximal displacement per step was set as 0.5 Å for translation and 1.25° for rotations and torsions.

After each randomly sampled move, the new ligand conformation is scored, as described by equation (1) with an additional term for the ligand internal energy, and this move is tested using a Metropolis criterion. The ligand internal energy is computed using the GAFF force field, as implemented in the OpenBabel API [38]. A graphical scheme of the MC procedure is shown in the Supplementary Material. For the MC simulations shown here, the MC temperature was set to 100K after preliminary tests.

The ensemble binding energy (ΔE_bind_) was computed by subtracting the energies computed for ligand within the receptor and the free ligand:

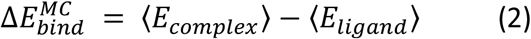

In addition to the energy terms, a conformational entropy (S_conf_) is estimated using a first-order approximation [39,40], where each degree of freedom for the ligand conformation is approximated to be independent of each other. In this scenario, for each degree of freedom, the entropy can be computed by computing the probability of the system in bins and then summing over the bins. The overall conformational entropy is given by the sum of the entropy for each degree of freedom. Again, the change in entropy is computed by the difference in the conformational entropy for the bound system and the free ligand. The final energy computed is described as:

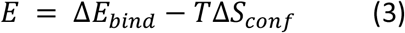

All MC simulations were run in five replicates using a different random seed for each simulation. The final binding energies are described as the average over the five simulations and the error in the binding energy is estimated as their standard deviation.

### 2.2 Protein Expression and Purification

PPARγ LBD was expressed and purified as previously described [41,42]. Briefly, hPPARγ-LBD was cloned into pET28a(+) plasmid and transformed in BL21 DE3 *Escherichia coli* strain. An initial was inoculated and grown at 37°C until the OD_600nm_ reached 0.8. Protein expression was induced with 1 mM isopropyl-β-D-thiogalactoside for 8 hours at 18°C. Cells were collected by centrifugation and disrupted by sonication, followed by centrifugation to remove cell debris. His-tagged protein was purified by metal affinity chromatography using a Talon Superflow Metal Affinity Resin by loading the supernatant into the pre-equilibrated resin. The protein was eluted with 300 mM imidazole in a buffer containing 20 mM TRIS-HCl, pH 8.0, 1 mM dithiothreitol, 100 mM NaCl and 5% glycerol. Purified protein was concentrated up to 10 mg/ml for crystallization assays.

### 2.3 Protein Crystallization and Structure Determination

Before protein crystallization, the protein solution was mixed in a 1:1 stoichiometry with ligand T2, keeping the DMSO concentrations under 5% in the final solution. A previously determined crystallization condition was used, where 1 μl of the protein was equilibrated against an equal volume of the reservoir solution containing 100 mM HEPES, pH 7.0 and 0.85 M sodium citrate. Suitable crystals grew after seven days at 18°C. Crystal was cryoprotected with 15% (v/v) ethylene glycol and rapidly cooled to 100K in a gaseous nitrogen stream prior to data collection in the MX-2 beamline of the LNLS [43]. Diffraction data were reduced using XDS [44] and scaled using AIMLESS program [45,46] from the CCP4 package [47]. The structure was solved by molecular replacement using PHASER [48]. Structure refinement was achieved by iterated cycles of manual building with COOT [49,50] and PHENIX.refine [51,52]. The final structure was validated using MOLPROBITY [53] as implemented in PHENIX. The final coordinates and structure factors were deposited in the PDB with a PDB ID 7LOT.

### 2.4 Competition Assay

To assess the ability of the screening ligands to bind PPARγ, the PolarScreen kit (Invitrogen) was used, following the manufacturer’s instructions. The ligands with ZINC IDs ZINC4999766, ZINC4999773 and ZINC4999768, in short T1, T2 and T3, were acquired from SPECS (catalog numbers AN-465/43369171, AN-465/43369504, AN-465/43369978) and were diluted in DMSO and incubated different concentrations with the receptor-fluormone complex. The displacement of the fluormone was measured by fluorescence polarization using an Envision plate reader (Perkin Elmer), with excitation/emission wavelengths of 485/535 nm. Before reading, the mixture was incubated for two hours at room temperature.

## 3. Results

### 3.1 Molecular docking suggests Tetrazole Compounds as a PPARγ ligand

An initial screening of PPARγ ligand candidates was set using a virtual screening strategy with the program DOCK 3.5 and, after a visual inspection of the results in UCSF Chimera [30], compounds that could make favorable interactions with the polar arm of PPARγ binding pocket were selected. Four tetrazole compounds were identified in this initial screening and, since the tetrazole moiety has not been yet described for PPARγ ligands, as far as these authors are aware, and considering that it is a bioisoster of the carboxylate group, usual among PPARγ agonists, we decided to further evaluate the potential of these compounds as PPARγ ligand candidates.

Three compounds, T1, T2 and T3, were selected for further investigations (Figure 1). In this analysis, the three compounds were re-docked using the program LiBELa [32–34]. This program uses a force field-based (AMBER FF14SB force field) interaction energy score in a combined, ligand- and receptor-based, search engine. These compounds have a tetrazole moiety docked on the polar region of the PPARγ ligand pocket composed of H323, H449 Y327, Y473, S289 and C285. An interesting hydrogen bond pattern could be visualized between residues Y327, H323 and S289 and the tetrazole moiety (A ring) while the B and C rings give a good fitting of the ligand into the ‘Y-shaped’ pocket, with structural similarity to rosiglitazone fitting on the receptor cavity [54], as shown in Figure 2.

**Figure 1.**
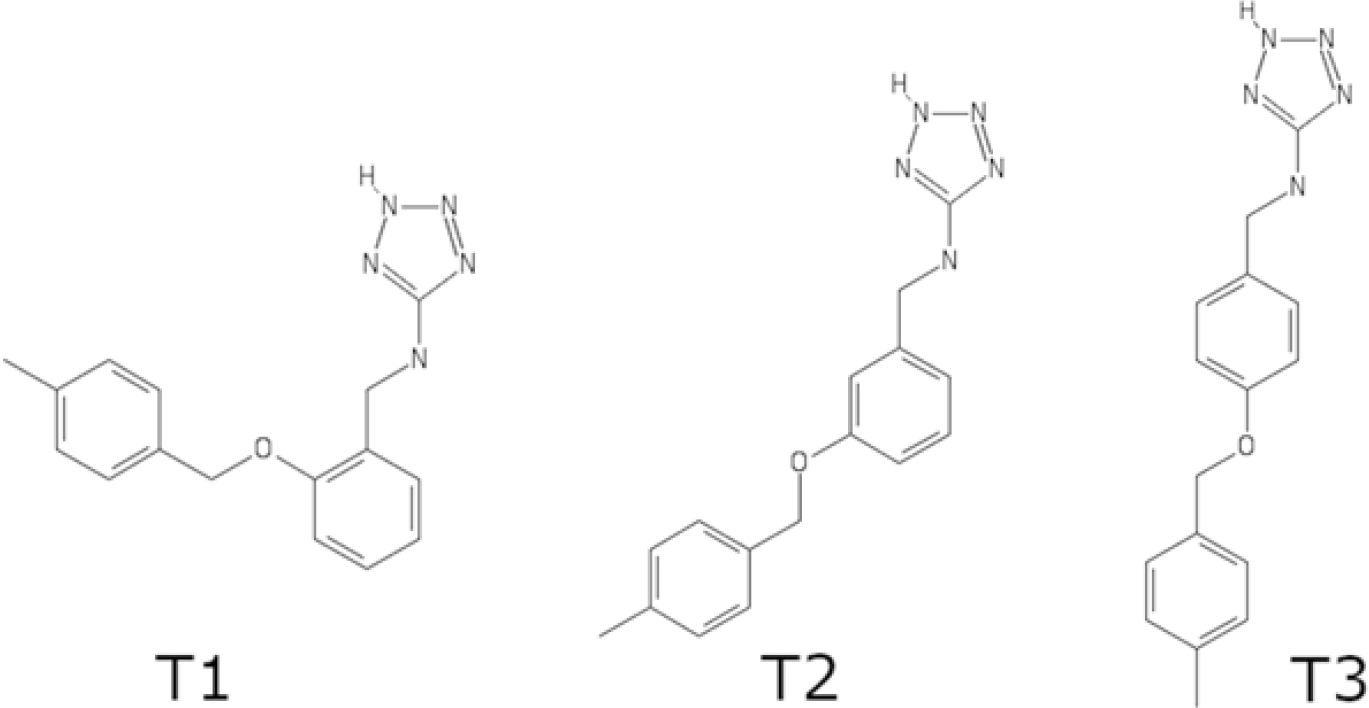
Structures of the tetrazole compounds T1, T2 and T3.

**Figure 2.**
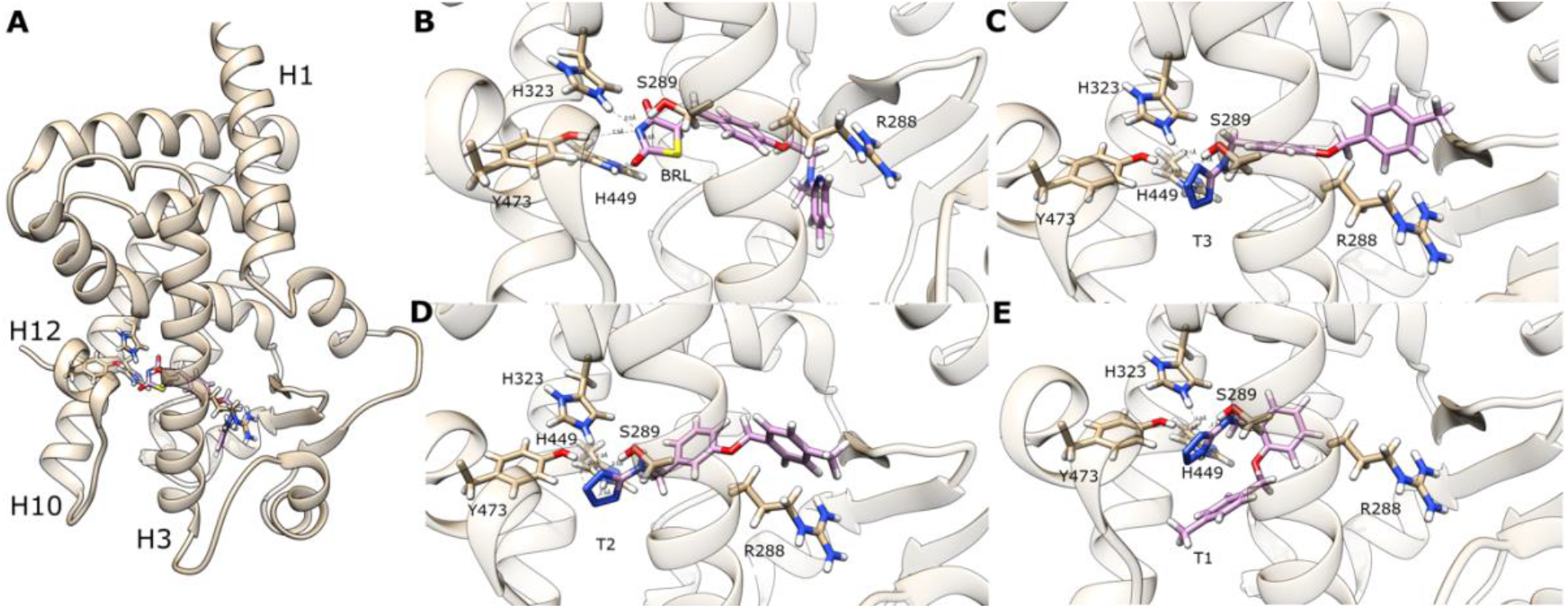
Binding conformations for rosiglitazone (BRL) and the compounds T1, T2 and T3 as predicted by docking calculations. (A) Overall view of the docked complex PPARγ-BRL. (B-D) Close view of residues of the polar arm involved in polar interactions with BRL, T3, T2 and T1, respectively.

The energy scores computed for these compounds with LIBELA using a PB-based electrostatic model and AMBER Lennard-Jones potential and a 10-12 hydrogen bond potential estimated the binding energy for rosiglitazone (BRL) as −58 kcal/mol, while the tetrazole compounds were ranked with interaction energies in the range between −61 and −55 kcal/mol, revealing reasonable binding poses as well as favorable interaction energies, comparable with the energies of the known high affinity ligand BRL.

The docked conformations for BRL, T1, T2 and T3 were used in Monte Carlo (MC) simulations of the ligand within the protein active site in LIBELA using the same energy potential as used in ligand docking. For this analysis, five independent MC simulations were run for each complex. The ligand coordinates were written every 1,000 accepted steps and the simulations were run for 50 million steps, resulting in 5,000 coordinates in a MC ‘*trajectory*’ for each simulation. The coordinates were clustered using the clustering tool as implemented in UCSF Chimera [30] and a ligand conformation of the most populated cluster is shown for each ligand in Figure 3.

**Figure 3.**
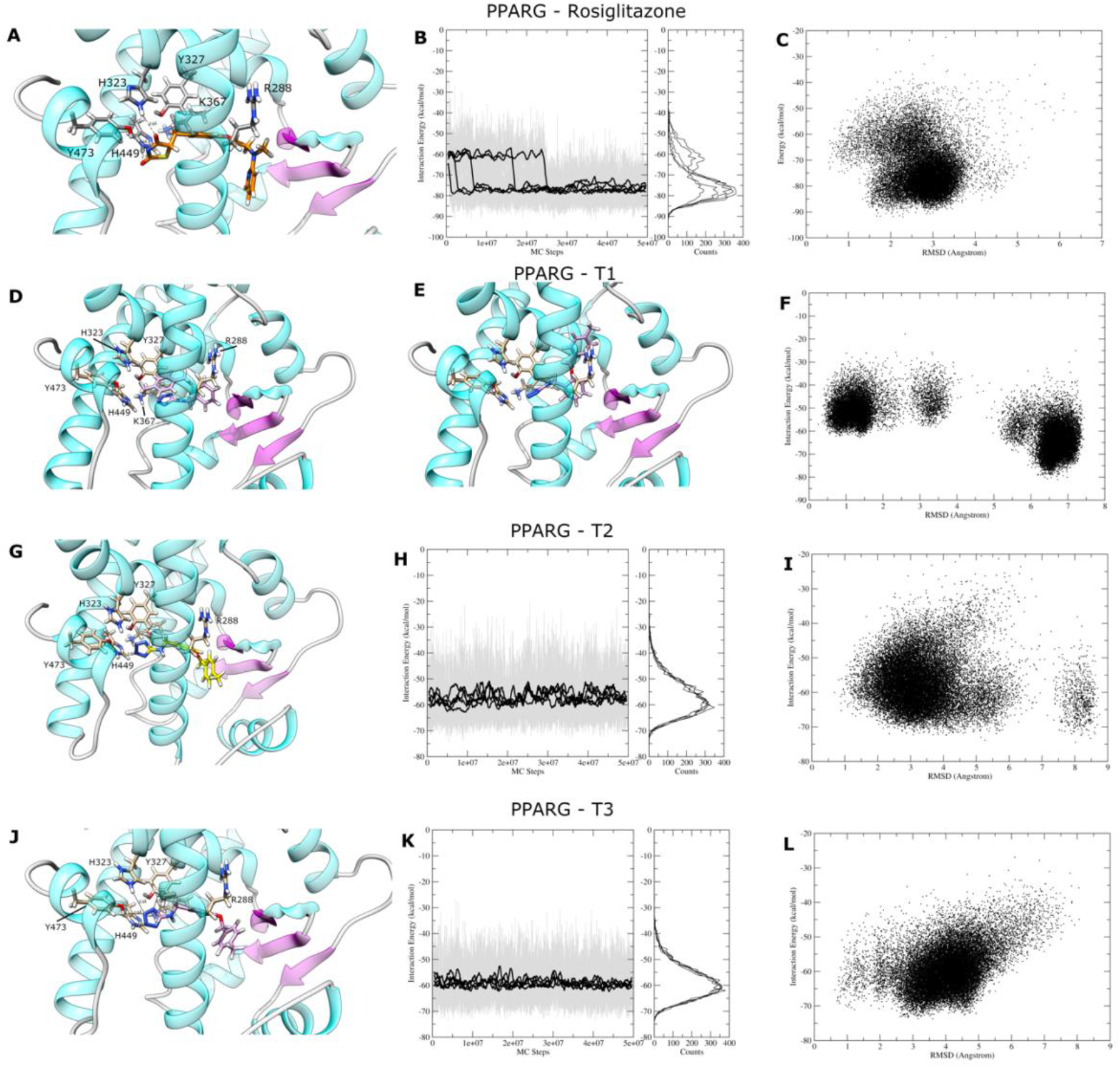
Analysis of the Monte Carlo simulations for BRL, T1, T2 and T3. A representative conformation of the most populated cluster in MC simulations for BRL is shown in panel A. The interaction energy per MC step is shown in the middle column for BRL, T2 and T3, together with a histogram analysis. The column in the right shows a scatter plot of the RMSD observed for the ligand during the simulation, as compared to the docked conformation, and its interaction energy. The two most populated conformations found for ligand T1 are shown in panels D and E.

As expected, the most populated cluster observed for rosiglitazone (BRL) is in close agreement with the experimental pose observed in the crystal structure (PDB ID 7AWC) solved at 1.7 Å, with hydrogen bonds observed for the TZD ring and H323, H449 and Y473, located in the helix H12 (Figure 3A). Eventual flips in the ring were observed during the simulation, but the experimental pose was the predominant conformation in the MC simulation, as indicated in the analysis of energy versus MC step (Figure 3B) and also in the analysis of energy versus RMSD (Figure 3C). For the later, the RMSD was computed using the docked conformation as the reference. An analysis of the interaction energies, computed as indicated in equation (2) and with the additional term for conformational entropy, indicated an interaction energy of −71 ± 3 kcal/mol for BRL. Here, the error was estimated by the standard deviation among the five independent simulations.

For ligand T1, two different ligand conformations were sampled during the MC simulations, resulting in two abundantly populated clusters in the analysis of the simulated coordinates. The first conformation (Figure 3D) shows the ligand curved within the binding pocket, with the tetrazole ring close to K367. In the second conformation (Figure 3E), the tetrazole ring is placed in the polar arm of the binding pocket, while the C ring points upward, close to R288. The analysis of the RMSD *versus* Energy (Figure 3F) also shows the two major clusters, with typical interaction energies around −50 kcal/mol and −65 kcal/mol. The analysis of the energies for the entire ensemble and taking into account the entropy loss resulted in an interaction energy of −49 ± 6 kcal/mol, thus significantly less favorable than the interaction energy observed for rosiglitazone.

For the ligands T2 and T3, similar binding poses were found, with the tetrazole ring docked in the polar arm and close to helix H12 and the B and C ring following a similar shape as observed for rosiglitazone (Figure 3G and 3J). In both cases, the simulations were dominated by a single *macrostate*, as shown in Figures 3I and 3L, and the analysis of the interaction energies revealed that T2 binds to PPARγ-LBD with an interaction energy of −57.1 ± 0.5 kcal/mol, while T3 has an interaction energy of −58.7 ± 0.2 kcal/mol.

The comparison of the energies computed with MC simulations and the docking energies shows that the sampling generated by Metropolis MC seems to correct or refine the energies computed with docking. The ranking of the ligands, after MC ensemble computation ranks rosiglitazone with an interaction energy of −71 kcal/mol, followed by T3 and T2, with −58 and −57 kcal/mol, respectively, and then T1 with −48 kcal/mol. Since rosiglitazone is a known high affinity PPARγ ligand, this ranking seems to be in line with what would be expected for screening campaigns. Similar strategies were explored with the combination of docking and molecular dynamic (MD) simulations [55–57] and, in most cases, the use of an ensemble generation *post-docking* results in a better ranking of the ligands. Here we show that a simplified approach, even based on the same underlying docking principles, seems to be applicable in this context.

It is also interesting to note the dependency of the ligand conformation in the binding profile. T1, T2 and T3 differ from each other by the *ortho, meta* and *para* substitutions in ring B (Figure 1). The MC simulations indicate that the *o-*substitution impairs the favorable filling of the Y-shaped binding pocket in PPARγ-LBD, while the *m-* and *p-*substitutions are both compatible and equally favorable.

### 3.2 Binding Data Indicates that T1, T2 and T3 are PPARγ Binders

The favorable interaction energies computed for the tetrazole compounds T1, T2 and T3 prompted us to evaluate their actual capacity to bind to the PPARγ ligand binding domain (LBD). Using the PPARγ PolarScreen Assay kit, preliminary competition curves were assayed for the three compounds. While rosiglitazone has an IC_50_ determined in this assay as 159 nM, the IC_50_ values for T1, T2 and T3 were determined to be in the range of 4 to 20 μM, with T1 showing a lower affinity (IC_50_ of 20 μM), and T2 and T3 with comparable and higher affinity, with an IC_50_ of 4 μM. Typical curves obtained in these experiments are shown in Supplementary Material. The ligand T2 was chosen for a deeper investigation and the binding curve for this ligand is shown in Figure 4 below. This data suggests that these tetrazole compounds are weak PPARγ binders, in the low micromolar range, which would be expected for compounds identified in a screening of compound libraries. It is interesting to note that the ligands found with higher and lower affinity are in line with the ranking as suggested by MC simulations shown above.

**Figure 4.**
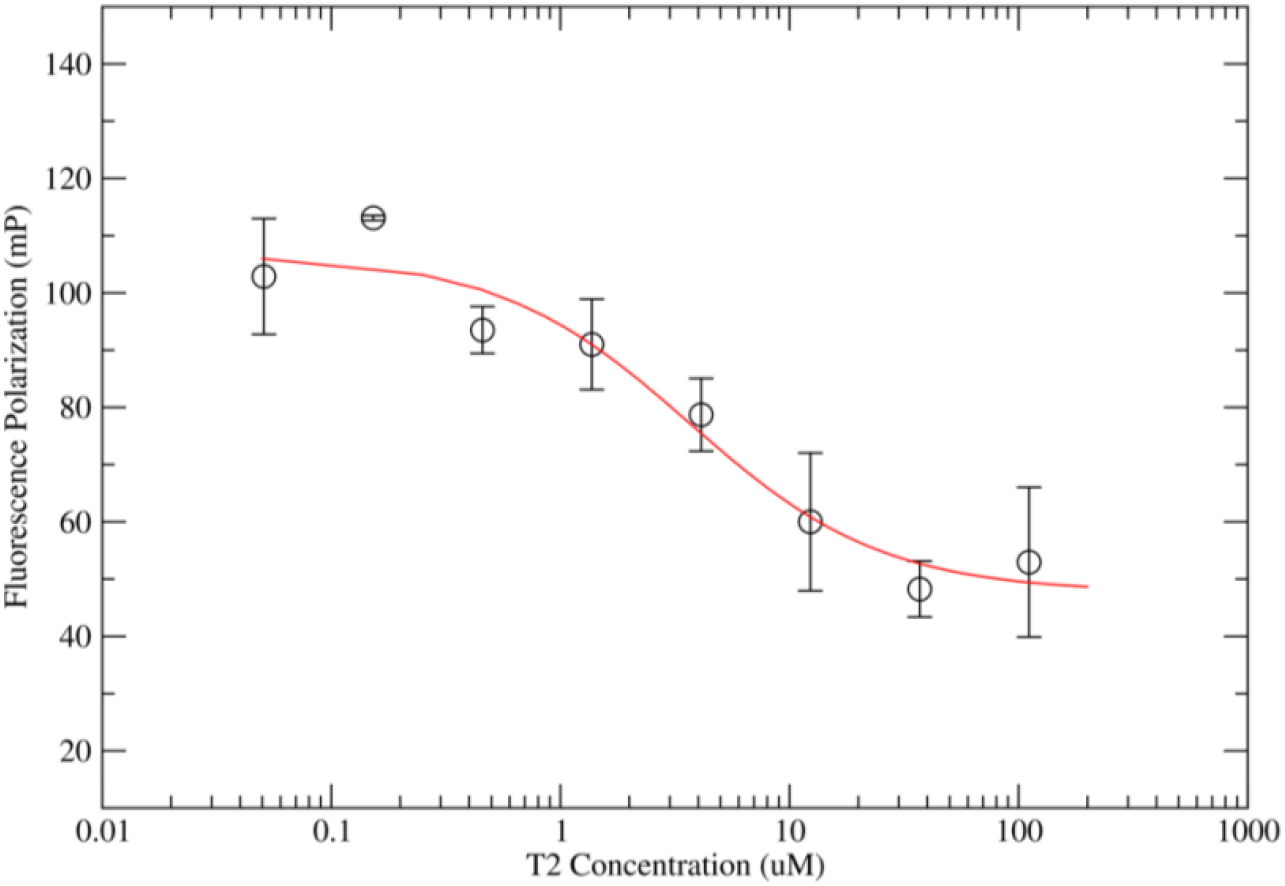
Binding curve obtained in PolarScreen assay for ligand T2. An IC_50_ of 3.7 μM was obtained by fitting the data in a ligand competition model.

### 3.3 Crystal structure of PPARγ-T2 complex

The investigation of the interaction between T2 and PPARγ was also deepened by an experimental investigation at an atomic level with the structural determination of this complex at a 2.3 Å resolution (Table 1). The crystal structure was composed of two molecules in the asymmetrical unit. As previously noted to this crystallization condition [58], one of the molecules has its helix 12 (H12) in an open conformation perfectly mimicking the co-activator interaction with a symmetrical molecule with H12 in a closed state. T2 appears in the two molecules, but with different binding modes, depending on the H12 conformation, suggesting an H12-dependent interaction and, possibly also an H12-dependent activation.

**Table 1.** Data collection and refinement statistics for PPAR gamma - T2 complex.

|  |  |
| --- | --- |
| PDB ID | 7LOT |
| Wavelength (Å) | 1.459 |
| Resolution range (Å) | 45.39 - 2.291 (2.373 - 2.291) |
| Space group | C 1 2 1 |
| Unit cell (Å,°) | 92.791 62.142 118.277 90 101.951 90 |
| Total reflections | 99747 (9935) |
| Unique reflections | 28560 (2745) |
| Multiplicity | 3.5 (3.6) |
| Completeness (%) | 93.30 (92.31) |
| Mean I/sigma(I) | 17.13 (2.08) |
| Wilson B-factor (Å<sup>2</sup>) | 54.12 |
| R-merge | 0.04274 (0.6472) |
| R-meas | 0.05043 (0.7608) |
| R-pim | 0.02657 (0.3979) |
| CC1/2 | 0.999 (0.822) |
| CC* | 1 (0.95) |
| Reflections used in refinement | 27880 (2738) |
| Reflections used for R-free | 1299 (144) |
| R-work | 0.2026 (0.2869) |
| R-free | 0.2593 (0.3343) |
| CC(work) | 0.960 (0.872) |
| CC(free) | 0.949 (0.825) |
| Number of non-hydrogen atoms | 4034 |
| macromolecules | 3947 |
| ligands | 44 |
| solvent | 43 |
| Protein residues | 512 |
| RMS(bonds) | 0.014 |
| RMS(angles) | 1.47 |
| Ramachandran favored (%) | 97.60 |
| Ramachandran allowed (%) | 2.40 |
| Ramachandran outliers (%) | 0.00 |
| Rotamer outliers (%) | 0.24 |
| Clashscore | 6.10 |
| Average B-factor | 74.39 |
| macromolecules | 74.38 |
| ligands | 88.79 |
| solvent | 60.03 |
| Number of TLS groups | 12 |

*Statistics for the highest-resolution shell are shown in parentheses.*

In chain A, where H12 is found in a ‘closed’ state, the best fit of the ligand into the density was found in a pose where the T2 polar head, the tetrazole group, contacts H449 (3.5 Å) and C285 (3.3 Å) (Figure 5). This conformation is very close to the conformation suggested by docking, highlighting the outstanding relevance of the H12 conformation for the docking pose, as we will show below. In chain B, where H12 is found in a ‘semi-opened conformation’, T2 is rotated and the tetrazole group is accommodated in the cleft among helix H3 and the β-strands. Interestingly, the C ring of the ligand makes cation-π interactions with R288 in this second conformation. Is it worth noting the potential stabilization of the β-strands in this conformation, which has a known effect on the receptor activation [59].

**Figure 5.**
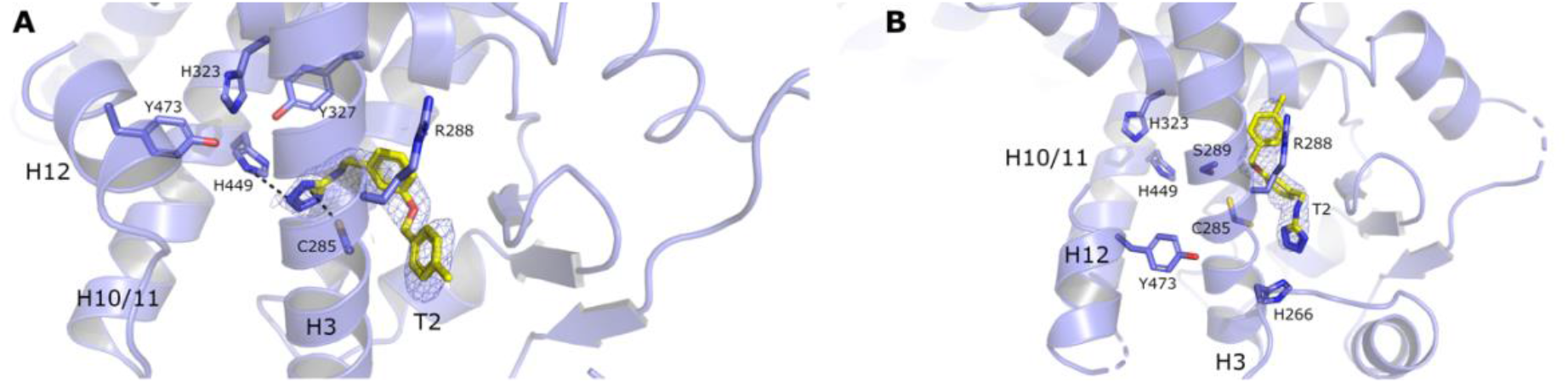
Crystal structure PPARγ bound to T2. (A) Binding conformation of T2 in the ‘A’ chain of PPARγ. In this chain, the H12 is found in a closed state. (B) Binding conformation for T2. In this chain, H12 is found in an open state. The electron density map (2F_obs_-F_c_) is shown in blue mesh contoured at 1σ.

The comparison of the conformations found in the opened and closed state of H12 suggests that the T2 can adopt two conformations depending on the conformation of H12. When the binding cleft is entirely formed, i.e., with Y473 contributing to create a polar cleft, the ligand interacts closely to H12. When H12 is not completely in a closed state (here, probably due to the crystal contacts that make H12 mimic a co-activator peptide) the binding cleft misses an important interaction residue, and this structural change seems to affect T2 binding. In this situation, the ligand can interact with R288. The ligand conformation found in the H12 opened stated resembles the conformation observed for ligands MRL-24 [12] and also for GQ-16 [60], where the ligand stabilizes the β-strands through some non-polar contacts. It is interesting to note that both MRL-24 and GQ-16 have been described as H12-independent partial agonists with decreased side effects.

## 4. Discussion

The recent reports from the regulatory agencies regarding the safety profile of the TZDs raised the question of whether PPARγ-mediated drugs are useful as insulin sensitizers. Recent findings, however, suggest that there are at least two main directions that deserve more investigation relating to the use of PPARγ ligands. The first line was firstly described by Spiegelman’s group in 2010 [59] when it was shown that a PPARγ partial agonist that activates the receptor through an H12-independent mechanism can block the phosphorylation of S275 by Cdk5 in adipose tissues. The phosphorylation of this residue is related to changes in the expression of several genes involved in obesity and insulin sensitization, including adiponectin. Thus, the development of small molecules able to bind to PPARγ without causing a full receptor activation but still blocking the phosphorylation may be useful as both insulin sensitizers and anti-obesity drugs.

A second direction involves the submaximal activation of the receptor. In this context, PAT5A, a PPARγ partial agonist, was shown to exhibit hypoglycemic effects with reduced adipogenic activity [61]. PAT5A was shown to partially activate PPARγ with reduced adipogenesis, which made the therapeutic profile more interesting. The same behavior was reported to 3-benzoyl or 3-benzisoxazoyl ligands [62]. Balaglitazone, another PPARγ partial agonist, also showed a safer profile as an antidiabetic agent, when compared to the full agonists TZD [23].

The discovery of tetrazole compounds as PPARγ binders is an academic effort in the second direction. Here we used a structure-based strategy to discover a ligand with different structural properties as a chemical probe to PPARγ active site binding and activation. Surprisingly, T2 revealed two different binding patterns, which seemed to be related to the H12 conformation. The closed conformation of H12 forms the classical binding pocket with a hydrogen bond involving H449 located in the polar arm of the receptor. On the other hand, when the H12 is not in the closed state, this network of hydrogen bonds is disturbed what seems to importantly alter the binding pattern, especially for partial agonists.

Bruning and coworkers showed already that some PPARγ partial agonists, such as MRL-24, for example, can activate the receptor without stabilizing the H12 [12]. This ligand binds to PPARγ subsite located among helix H3 and the β-strands, as shown by H/D exchange. Waku and coworkers also showed that the anti-inflammatory indomethacin (IDM) is a PPARγ partial agonist [63]. The crystal structure of IDM bound to PPARγ unexpectedly showed two ligand molecules occupying the active site in two subsites. The first subsite was located in the canonical PPARγ site (AF2 subsite), where the ligand interacted with Y473 in the H12, H449, and H323, the classical residues involved in full agonists binding. On the other hand, a second subsite was identified close to H3 and the β-strands, where many hydrophobic interactions stabilize the region of the receptor known to directly interact with RXR DBD [4]. The same behavior was observed for carboxamides bound to PPARγ [64].

Taken together, the binding modes observed to date for PPARγ ligands can be grouped in two structurally distinct patterns. The first pattern involves polar interaction among polar groups (usually acting as acceptors in a hydrogen bond) of the ligand and helix H12 (Y473), together with the polar cluster that forms the arm I of the receptor binding pocket (H449, Y327, and H323). Many of these ligands have a high affinity for the receptor and promote full receptor activation. The second pattern of ligand interaction is observed with MRL-24 and GQ-16. These ligands have little or no polar interaction with the polar arm of the binding pocket and act mainly through the stabilization of the β-strands by non-polar interactions. The tetrazole compound T2 described here shows an intermediate binding pattern where some hydrophobic interactions do stabilize the β-strand region of the ligand binding domain while a single polar interaction contributes to a minimum H12 stabilization. A similar pattern was observed previously in the interaction of INT131 with PPARγ [65]. This ligand also contacts PPARγ H12 through a water-mediated interaction and, interestingly, is still able to activate the receptor in the presence of a mutation that replaces Y473 for an alanine. Finally, this partial agonism observed for INT131 resulted in serum glucose reduction without edema and other side effects [66–68], resulting in increased therapeutic safety.

Regarding the tetrazole compounds shown here, a still open question is whether the tetrazole ring is responsible for the decrease in the affinity for T2, as compared to rosiglitazone, and causing a potential partial agonism effect. To address this question, we replaced the TZD ring in rosiglitazone with a tetrazole ring creating a new ligand that morphs the structural properties of T2 and rosiglitazone. After an MC simulation following the same protocol as used for other ligands, the virtual compound showed an interaction energy of −61.4 ± 0.9 kcal/mol, i.e., an interaction energy significantly higher than the interaction observed for rosiglitazone (−71 kcal/mol) and closer to the T2 interaction energy (−57 kcal/mol). This result suggests that the interactions between the tetrazole ring and the receptor LBD are the major reason for the lower affinity and, possibly, for the partial agonism of the tetrazole compounds.

Do the tetrazole compounds identified here have any relevant effect *in vivo*? Although *in vivo* studies were not accessible to us, the hypoglycemic effects of perfluoro-N-[4-(lH-tetrazol-5-ylmethyl)phenyl]-alkanamides, tetrazoles somewhat similar to T2, were described previously [69,70], with good pharmacokinetic profile [71]. When tested in *ob*/*ob* mice in single oral doses for two consecutive days some tetrazoles resulted in a hypoglycemic effect similar to the effect observed for ciglitazone, although at higher doses [69]. This experimental evidence of the hypoglycemic effect of tetrazole molecules similar to T2 makes it tempting to speculate that T2 may have a hypoglycemic effect *in vivo*, with partial agonism in PPARγ, as expected from the binding mode.

In conclusion, here we showed the screening and structure-based identification of tetrazole compounds as novel PPARγ ligands. The crystal structure of the PPARγ-T2 complex was determined confirming the binding mode. Besides, Monte Carlo simulations revealed the most important interactions between the tetrazole ring and the polar arm (arm I) within the ligand binding pocket.

## Acknowledgements

This research used resources of the Brazilian Synchrotron Light Laboratory (LNLS). The MX2 beamline staff is acknowledged for the assistance during the experiments, as well as the LNBIO staff. The authors also thank the financial support from the Fundação de Apoio À Pesquisa do Estado de São Paulo (FAPESP) through grants 2010/15376-8, 2014/06565-2, 2015/26722-8 and 2020/03983-9 and from the Conselho Nacional de Desenvolvimento Científico e Tecnológico (CNPq) through grants 303165/2018-9, 485950/2013-8 and 476606/2010-1. This study was also financed in part by the Coordenação de Aperfeiçoamento de Pessoal de Nível Superior - Brasil (CAPES) - Finance Code 001.

