## Supplemental Figures S1 and S2 for "Tetrazoles as PPARγ ligands: A Structural and Computational Investigation"

**Supplementary Figure 1**. Typical binding curves for ligands T1 and T3 in the PolarScreen assay.

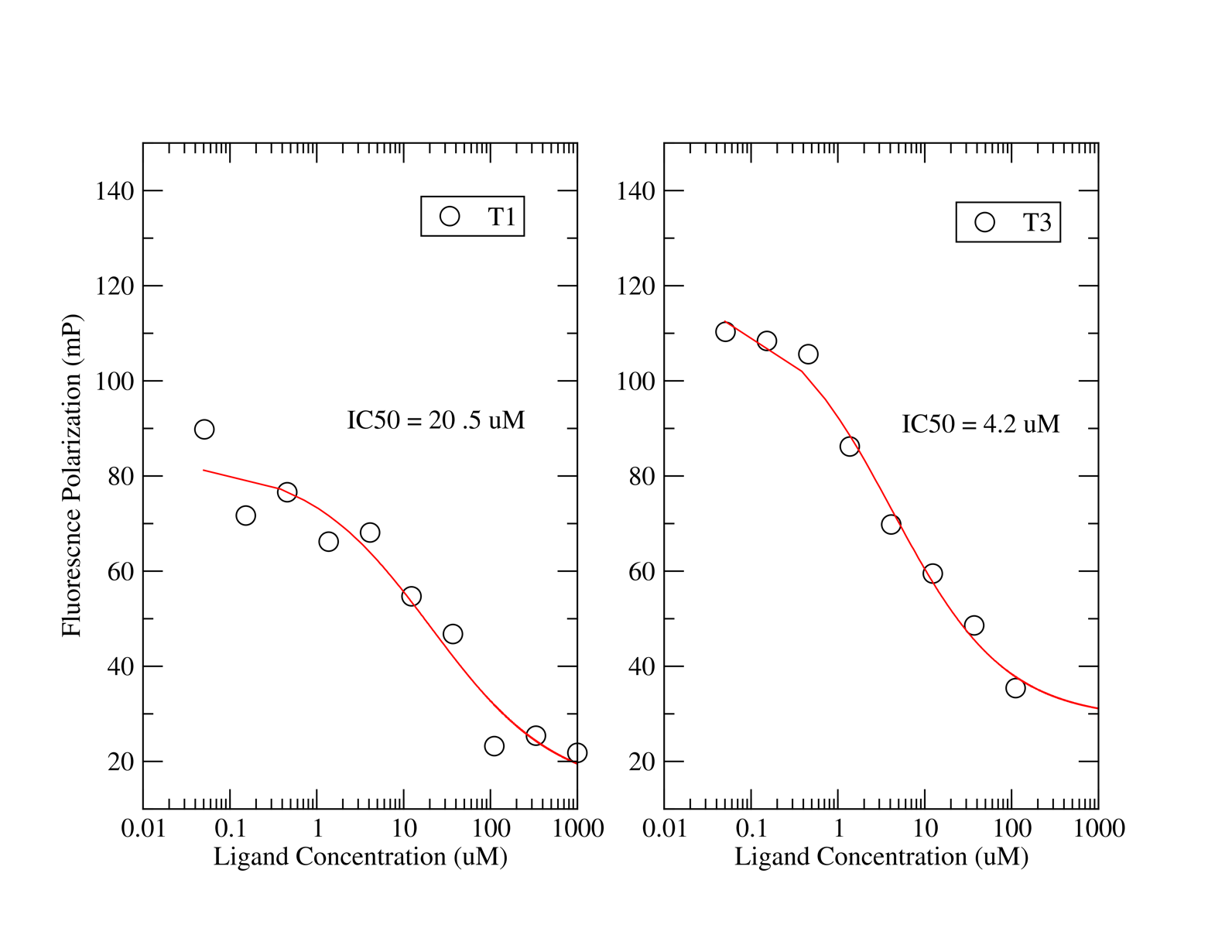
